# Metagenomic sequencing to replace semi-quantitative urine culture for detection of urinary tract infections: a proof of concept

**DOI:** 10.1101/178178

**Authors:** Victoria A. Janes, Sébastien Matamoros, Niels Willemse, Caroline E. Visser, Bob de Wever, Marja E. Jakobs, Poorani Subramanian, Nur A. Hasan, Rita R. Colwell, Menno D. de Jong, Constance Schultsz

**Affiliations:** Department of Medical Microbiology, Academic Medical Center, University of Amsterdam, Amsterdam, The Netherlands.; CosmosID, Rockville, Maryland, United States of America.; Department of Global Health – Amsterdam Institute for Global Health and Development, Amsterdam, The Netherlands.

**Author notes:** Corresponding author: Victoria A. Janes.

## Abstract

Semi-quantitative bacterial culture is the standard method to diagnose urinary tract infections (UTI), but bacterial growth rate limits diagnostic speed and it is unreliable when patients have been pre-treated with antibiotics. Metagenomics could increase diagnostic speed and accuracy by sequencing the microbiome and resistome directly from urine samples, bypassing culture. However, a semi-quantitative approach – as needed for diagnosing UTIs – has not been established.

Metagenomics was deployed to identify and semi-quantify bacterial presence indicative of UTI, predict antimicrobial susceptibility (AMR), and results were compared to semi-quantitative culture. Whole genome sequencing of the corresponding uropathogens was done for comparison. Analysis time and cost were tracked.

Forty-one consecutive urine samples underwent metagenomic analysis. All culture positive samples contained >200ng of DNA, suggestive of a threshold below which UTI could be ruled out solely based on DNA quantity. A semi-quantitative Diagnostic Index (DI) was created by multiplying the total DNA quantity by the relative abundance of uropathogens per urine sample. The DI allowed discrimination of UTI from non-UTI samples in all but 1 case. Metagenomic detection of AMR determinants correctly predicted the phenotype of uropathogens in 20 of 32 cases. The metagenomic work-flow was 31h and cost €116 per sample, but could be reduced to 4.5h and €5 for low-DNA-yield non-UTI samples.

The genomic determinants of AMR and their distribution across uropathogens need to be better understood for prediction of AMR phenotypes by metagenomics. The introduction of the DI demonstrates the potential of semi-quantitative metagenomics to replace culture as rapid diagnostic method for UTI.

## INTRODUCTION

Urinary tract infections (UTIs) are amongst the most common infections that require antibiotic prescription, and in catheterized and hospitalized patients UTIs are the most common nosocomial infections(1-3). The high incidence of UTIs, paired with increasing resistance to first and second line antibiotics for common uropathogens, such as *Escherichia coli*, stress the need for rapid diagnostics in aid of appropriate antimicrobial treatment(4). However, the current standard for diagnosing UTIs is time-consuming as it is based on (semi-)quantitative culture of urine, followed by identification and antimicrobial susceptibility testing (AST) of isolated pathogens, which usually takes 1-3 days and for some pathogens up to 7 days. In addition, growth of clinically relevant bacteria may be hampered by prior antimicrobial treatment, which is common practice worldwide, resulting in diagnostic challenges and risks of misdiagnoses.

Dependence on culture for UTI diagnosis can be bypassed by quantitative molecular detection of uropathogens in urine specimens. Multiplex real-time PCR-based methods have shown promise for this purpose but are hampered by exclusive detection of selected micro-organisms and lack of, or incomplete information on antimicrobial resistance (AMR) patterns(5, 6). Direct metagenomic sequencing of microbial communities in urine provides opportunities for unbiased detection of uropathogens, including the presence of AMR genes, while current next generation sequencing (NGS) methods and bio-informatic tools facilitate generation of such results in a timely fashion. Metagenomic approaches have indeed shown promise in identifying uropathogens and their antimicrobial susceptibility patterns in urine samples(7). However, semi-quantitative detection of uropathogens is essential in order to replace culture-based diagnostics by metagenomic approaches since international clinical guidelines use bacterial loads (e.g. < 10^3^ or 10^5^ CFU/mL) to define evidence of UTI in different patient populations(8, 9). To facilitate such semi-quantitative metagenomic diagnostics for UTI, we developed and tested a metagenomic sequencing-based algorithm for determination of clinically relevant levels of bacterial DNA, indicative of urinary tract infections. Results were compared to the reference test semi-quantitative urine culture followed by species identification by MALDI-TOF, and automated AST. Whole genome sequencing (WGS) of isolated uropathogens served as benchmark for expected AMR gene content in metagenomes. We recorded the time to result and direct costs. The aim of this study was to provide proof of concept for the use of metagenomics as a tool for detecting UTI and predicting antimicrobial susceptibility patterns.

## METHODS

### Urine Samples

During two days, all urine samples obtained from patients with suspected UTIs that were assessed at the clinical microbiology laboratory for culture, identification, and AST, were included in the study. We collected up to a maximum of 20ml of surplus urine. Urine samples, corresponding routine culture plates and AST results were collected and anonymized, according to local and national ethical requirements for diagnostic studies using surplus routine care patient samples.

### Urine Culture

Urine was cultured according to local standard operating procedures (see supplementary data for summarized SOP). Bacterial isolates were identified morphologically and by MALDI-TOF MS (MALDI Biotyper, Bruker, Karlsruhe, Germany) and were reported in a semi-quantified way as the number of colony forming units (CFU) per ml.

A culture-positive urine was defined as growth of clinically relevant bacteria at numbers >10^3^ CFU/ml and for the purpose of this study all culture-positive urine samples were assigned to the UTI group. Growth of 10^3^ CFU/ml or less was reported as “no significant growth”, while mixed growth of commensal vaginal, rectal, or mucocutaneous bacteria in any numbers or absence of any growth were reported as “commensal flora” or “no growth”, respectively. The latter three culture results together formed the non-UTI group. The clinically relevant bacteria from culture-positive urines were subcultured on Columbia agar plates with 5% sheep blood (Biomérieux, Marcy-l’Étoile, France) prior to DNA extraction and WGS.

### Antimicrobial susceptibility testing

AST was done directly on urine by disc diffusion if Gram staining of the urine revealed the presence of abundant bacteria of a single morphology, or on identified bacterial isolates using VITEK 2 (version 06.01; Biomérieux), both according to EUCAST guidelines and breakpoints(see Supplementary data, Summary of SOPs)(10).

### DNA extraction and sequencing

DNA was extracted from 20 ml of urine. If <20 ml of urine was available, the sample was supplemented with PBS. The urine was first centrifuged for removal of human cells (2000 g, 30 s), followed by centrifugation of the supernatant for pelleting of bacterial cells (8000 g, 10 min). The bacterial pellet was pre-lysed with an in-house enzyme cocktail comprising achromopeptidase, mutanolysine, lysostaphine (1000:100:3) and lysozyme (1mg/ml) (Sigma-Aldrich, St. Louis, MO, USA) in TE buffer. This was followed by lysis using proteinase K and an in-house lysis buffer (Sodium-docecyl-Sulphate (1 %), Tween-20 (0.5 %) and Sarkosyl (0.5 %) in TE-buffer), after which automated DNA extraction was performed immediately using the NucliSENS easyMag platform (Biomérieux) following manufacturer’s instructions. DNA from the cultured bacteria was extracted with the Wizard Genomic DNA Purification Kit (Promega, Madison, Wisconsin, USA). The Qbit dsDNA HS Assay Kit (ThermoFisher, Waltham, Massachusetts, USA) was used to measure DNA concentration. The Ion Xpress^™^ Plus Fragment Library Kit (Thermo Fisher Scientific) was used for PCR-free, manual library preparation according to manufacturer’s specifications. Library quantification was performed with the Ion Library TaqMan Quantification Kit (Thermo Fisher Scientific). DNA was sequenced on the Ion Torrent Proton platform, set to produce an average of 1 million 200 base-pair length single-end reads per sample. Trimmomatic was used to remove low quality reads (settings WGS: headcrop 15, sliding window 4:20, Phred score 15; settings metagenomics: headcrop 15, crop 270, sliding window 4:20, Phred score 10, minlen 50)(11).

### Metagenomic analysis

Human sequences in the metagenomic datasets were removed with Deconseq version 0.4.3, human genome version 37(12). The remaining reads were analyzed for identification of microbial DNA at subspecies level and determination of the organism’s relative abundance using the CosmosID bioinformatics software package (CosmosID Inc., Rockville, MD)(13-16). A cloud version is accessible at https://app.cosmosid.com. The relative abundance of each bacterial organism per sample was expressed as a percentage of the total number of bacterial reads belonging to that organism, normalized for organism-specific genome length. Reads identified as eukaryote, viral, or archaeal were excluded.

All bacteria identified by metagenomics were classified as commensal or uropathogen. Commensal bacteria comprised members of the genito-urinary tract and skin microbiota generally not considered as urinary pathogenic. Uropathogens comprised all *Enterobacteriaciae,* other species known as common causative agents of UTI and putative uropathogens that have been described to cause UTI in rare cases(17). A full list of bacterial species detected and their designation is provided in the Supplementary data, table 1.

**Table 1.** Comparing species identification by semi-quantitative culture and metagenomics' relative abundance (Rel. ab.) of organisms per urine sample of the UTI group.

| Sample ID | Culture |  | Metagenomics |  |
| --- | --- | --- | --- | --- |
|  | Species | CFU/ml | Species | Rel. ab. (%) |
| 7 | *Enterobacter cloacae* | $10^4$ | *Enterobacter cloacae* | 58.8 |
|  |  |  | *Enterobacter species* | 34.8 |
|  |  |  | *Bacillus cereus* | 6.4 |
| 8 | *Staphylococcus aureus* | $>10^4$ | *Staphylococcus aureus* | 76.3 |
|  |  |  | *Klebsiella species* | 23.7 |
| 18 | *Escherichia coli* and skin flora | $>10^4$ | *Escherichia coli* | 55.8 |
|  |  |  | *Bifidobacterium pseudocatenulatum* | 25.0 |
|  |  |  | *Bifidobacterium longum* | 11.4 |
|  |  |  | *Shigella flexneri* | 3.1 |
|  |  |  | Other | 4.7 |
| 19 | *Escherichia coli* | $>10^5$ | *Klebsiella species* | 90.4 |
| | *Klebsiella pneumoniae* | $>10^5$ | *Klebsiella pneumoniae* | 8.1 |
|  |  |  | *Escherichia coli* | 0.9 |
|  |  |  | Other | 0.7 |
| 23 | *Escherichia coli* | $10^4$ | *Escherichia coli* | 100.0 |
| 28 | *Escherichia coli* | $10^4$ | *Escherichia coli* | 86.6 |
|  |  |  | *Shigella sonnei* | 6.6 |
|  |  |  | Other | 6.8 |
| 30 | *Escherichia coli* | $>10^5$ | *Escherichia coli* | 80.4 |
| | *Staphylococcus aureus* | $10^4$ | *Anaerococcus lactolyticus* | 9.3 |
|  |  |  | *Escherichia species* | 5.3 |
|  |  |  | *Peptoniphilus harei* | 3.6 |
|  |  |  | *Staphylococcus aureus* | 0.0 |
|  |  |  | Other | 1.5 |
| 45 | *Proteus mirabilis* | $>10^5$ | *Proteus mirabilis* | 66.4 |
| | *Morganella morganii* | $>10^5$ | *Morganella morganii* | 26.9 |
|  |  |  | *Klebsiella variicola* | 2.1 |
|  |  |  | *Klebsiella pneumoniae* | 1.7 |
|  |  |  | Other | 2.8 |
| 49 | *Enterobacter cloacae* | $>10^4$ | *Enterobacter cloacae* | 53.9 |
| | *Pseudomonas aeruginosa* | $>10^4$ | *Escherichia coli* | 29.4 |
|  |  |  | *Atopobium vaginae* | 8.9 |
|  |  |  | *Pseudomonas aeruginosa* | 7.8 |

Finally, the resistome i.e., the pool of resistance genes present in the microbial community, was characterized from the metagenome with the CosmosID (CosmosID Inc., Rockville, MD) metagenomic software package using CosmosID’s curated antibiotic resistance gene database. Resistome predictions were compared with results of AST and WGS of the corresponding isolated bacteria.

### Whole genome sequencing

SPAdes/3.6.0 was used for read-assembly (settings: --iontorrent, -k 21,33,55 -- careful)(18). KmerFinder version 2.1 identified the bacterial species(19). If KmerFinder was inconclusive, a BLAST search of the NCBI database was done, using the assembled genome(20).

ResFinder version 2.1 was used with default settings (90% identity match, 60% coverage) for detection of acquired antimicrobial resistance (AMR) genes (https://cge.cbs.dtu.dk/services/ResFinder/ analysis date: 14^th^ June 2017)(21). Additional acquired and chromosomal genes and mutations were detected from the assembled reads using the web-based “Resistance Gene Identifier” tool for searching the Comprehensive Antibiotic Resistance Database (CARD) using “strict” and “perfect” matches only with identity match of ≥99%(https://card.mcmaster.ca/analyze/rgi analysis date: 14^th^ June 2017)(22). AMR genes identified by WGS served as benchmark of expected AMR gene content in metagenomic sequence data.

### Metagenomic quantification

In order to integrate both DNA quantity and relative abundance in a single diagnostic measure, thus emulating the qualitative and quantitative properties of the reference test culture, we created the diagnostic index (DI). The DI is the product of the total quantity of extracted DNA in nanograms (D) and the relative abundance of uropathogens (RA) for each urine sample: DI = D * RA. Semi-quantitative culture and metagenomics results were compared one on one per urine sample for species identification and quantification. The median and interquartile ranges (IQR) for DNA yield, RA and DI for the groups UTI and non-UTI were compared. Thus, we assessed which metagenomic measure had best discriminatory power, displaying the least overlapping values between groups UTI and non-UTI.

### Statistics

Differences in relative abundance between uropathogens, as well as differences in total DNA yield between specimens were analyzed using the Mann-Whitney-U test (RStudio version 0.99.902). A significant difference was defined as a p-value <0.05. Graphs were designed with RStudio version 0.99.902. Sensitivity, specificity, and 95% confidence intervals (95% CI) were calculated using MEDCALC (https://www.medcalc.org/calc/diagnostic_test.php).

## RESULTS

### Culture and WGS of isolates

Forty-six consecutive urine samples collected for routine culture were included, results of which were reported as no-, commensal- or no significant (≤ 10^3^ CFU/ml) growth for respectively 12, 19 and 4 urines, hence classified as the non-UTI group. Eleven samples were classified as culture-positive and constituted the UTI-group. From these 11 specimens, 15 uropathogens were identified in total: 2 *Enterobacter cloacae,* 6 *Escherichia coli,* 2 *Klebsiella pneumoniae,* 2 *Staphylococcus aureus* and 1 each of *Proteus mirabilis, Morganella morganii,* and *Pseudomonas aeruginosa.* Of these, 14 were available for WGS. Species identification by WGS confirmed identification by MALDI-TOF in all 14 cases.

### Metagenomics of urine samples

Metagenomic sequencing and analysis was performed for 41 of the 46 included urine samples. In the UTI group, one sample did not contain surplus urine for metagenomic analysis, and DNA extraction was unsuccessful for another due to technical difficulties. In the non-UTI group, zero DNA was extracted from three samples for which culture showed no growth for one and commensal flora in 10^2^CFU/ml for two samples. Thus, 9 UTI and 32 non-UTI urine samples, containing 5.7-1210.0 ng of DNA, were used for metagenomic sequencing and analysis.

### Non-UTI group

The 11 sequenced urine samples yielding no growth in culture, showed at least one bacterial species to be present by metagenomics. In these samples, the most frequently detected genera were *Gardnerella, Bifidobacterium, Enterococcus,* and *Lactobacillus* (Supplementary data table 2-C). Of the 19 urine samples that grew commensal flora in culture, 17 were analyzed by metagenomics and the most frequently observed bacterial taxa were *Bifidobacterium, Lactobacillus, Prevotella*, *E. coli, Staphylococcus epidermidis,* and *Gardnerella vaginalis* (Supplementary data table 2-B). All four samples with no significant growth in culture, respectively showed *E. coli, Bifidobacterium* and *Lactobacillus* as most abundant taxa (Supplementary data table 2-D).

**Table 2.**
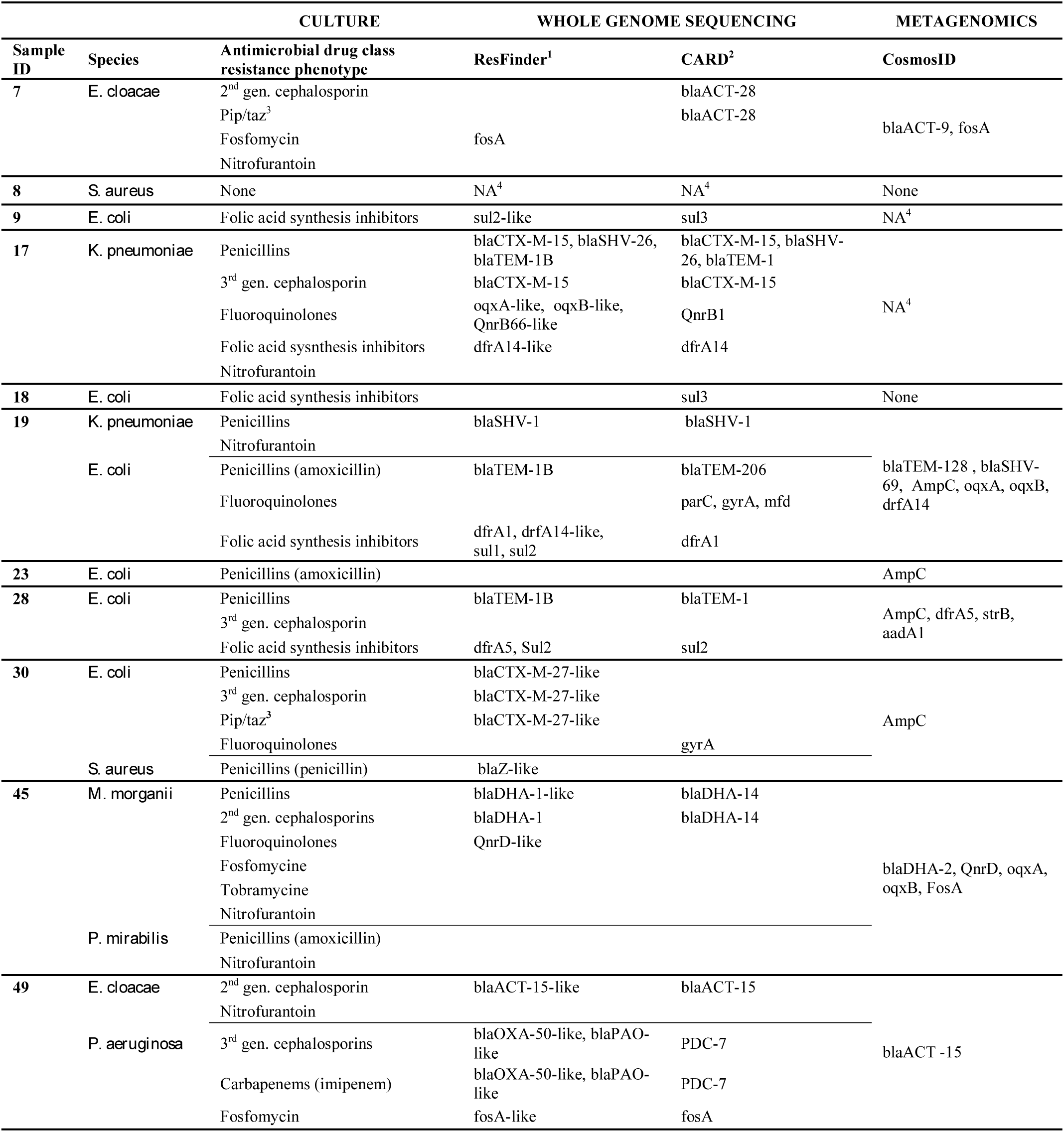
Phenotypic AST per drug class compared to genotypic AMR gene detection using the detection tools ResFinder and CARD (WGS), and CosmosID (metagenomics). Legend Table 2. If only 1 antibiotic was tested per antimicrobial drug class, the tested drug was noted between brackets. **^1^** Database contains acquired AMR determining genes only. **^2^** Database contains chromosomal and acquired AMR determining genes and mutations. **^3^**Piperacillin/tazobactam. **^4^** Not available.

### UTI group

In 9 of 11 urine samples from the UTI group that underwent metagenomic sequencing and analysis, 12 of the 13 cultured uropathogens were identified amongst the most abundant species by metagenomics, but some differences were observed (Table 1). Where *S. aureus* (sample 30), present in mixed growth with *E. coli*, was cultured at a concentration of 10^4^CFU/ml, it was not found in the metagenomic dataset. Sample 19 showed *Klebsiella pneumoniae* dominance (90.4% relative abundance) with only 0.9% relative abundance of *E. coli*, whilst culture suggested equal growth of >10^5^CFU/ml of both species. Similarly, culture of sample 49 suggested equal growth of *Enterobacter cloacae* and *Pseudomonas aeruginosa*, whilst metagenomics revealed *E. cloacae* dominance (relative abundance 53.9%) with *P. aeruginosa* present at 7.8% relative abundance. Additionally *E. coli* was detected at a RA of 29.4%. Only *S. aureus* (10^4^ CFU/ml) was cultured from sample 8 whilst *Klebsiella species* was additionally detected by metagenomics (Table 1) in a patient pre-treated with trimethoprim. Metagenomics identified additional taxa compared to culture in 6 other UTI samples including *Bifidobacterium species, Shigella species, Lactobacillus species, Bacillus cereus, Anaerococcus lactolyticus, Peptoniphilus harei* and *Atopobium vaginae.*

### Metagenomic quantification

The median relative abundance (RA) of uropathogens in urine samples from the UTI-group (93.6%; IQR 7.4) was significantly different (p<0.001) from that of the non-UTI group (0.3%; IQR 11.2) (Supplementary data, figure 1). The median DNA yield from urine samples of the UTI-group (642.4ng; IQR 456.5; range 203.5 – 1210.0ng) was and significantly different (p<0.001) from the non-UTI group (127.3ng; IQR 295.3; range 0.0-734.8 ng)(Supplementary data, Figure 2). Where the non-UTI samples had a variable DNA yield, all urine samples from the UTI group, yielded > 203 ng of DNA, forming a clear threshold below which no UTIs were found. However, both RA and DNA yield lacked discriminatory power as individual sample values were overlapping (Supplementary Data, figures 1 and 2).

**Figure 1.**
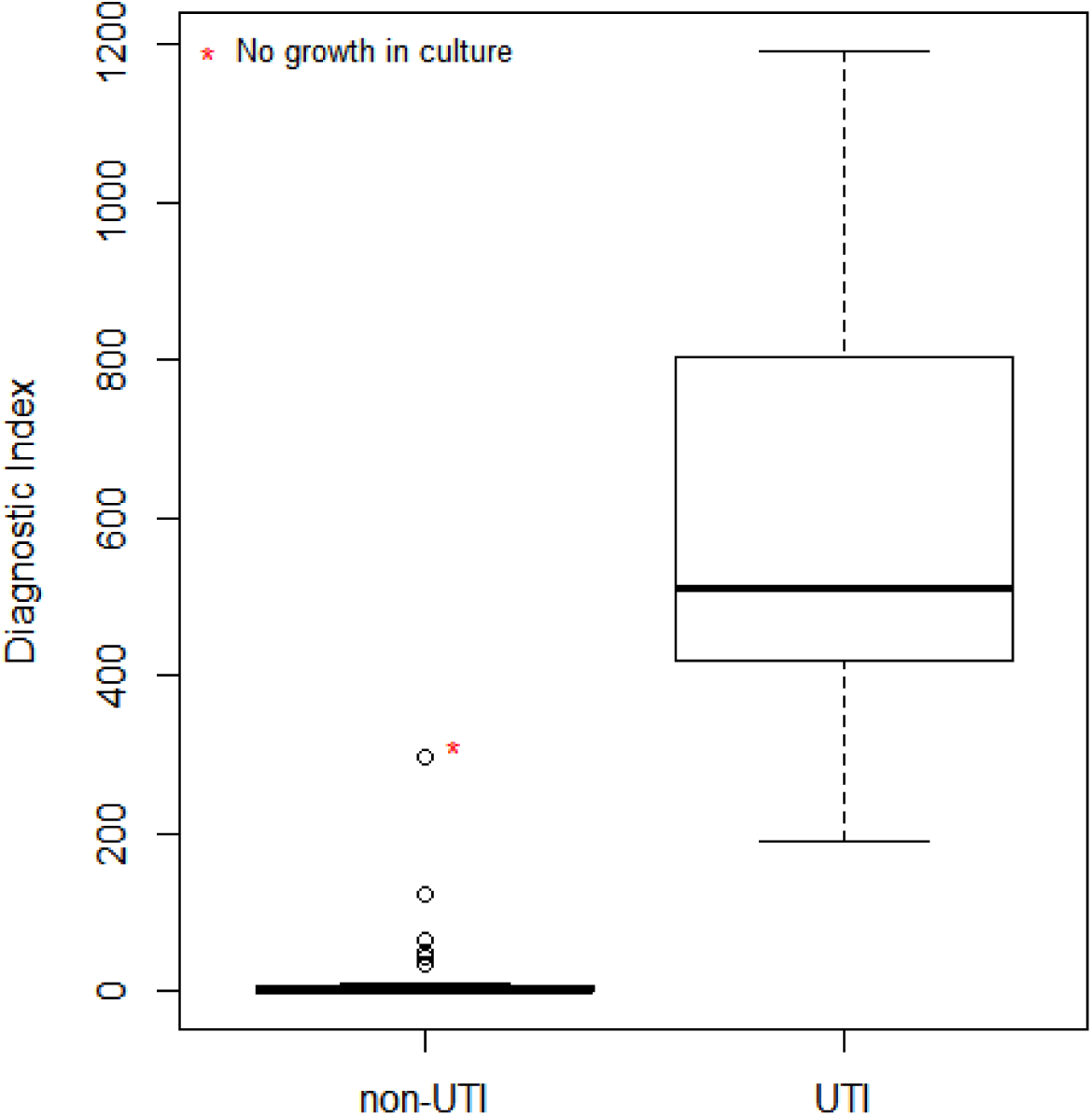
Diagnostic Index (DI) for urine samples in the groups UTI and non-UTI. Legend Figure 1. The “Diagnostic Index” (DI), the product of the relative abundance of uropathogens, and total DNA quantity per sample was computed for the groups UTI and non-UTI. The median DI for the UTI group was 512.0 (IQR 382.8) versus 0.11 (IQR 5.4) for the non-UTI group, which was a significant difference (p < 0.00001, Mann Whitney-U test). The red asterisk represents urine sample 44 showing *Aerococcus urinae* in metagenomic analysis (88% relative abundance, 325.6ng DNA extracted from that sample), which was classified as a uropathogen in this study. Culture showed no growth.

Integrating RA and DNA yield per urine sample in the diagnostic index (DI) discriminated all 9 culture positive urine samples of the UTI-group from the remaining 32 non-UTI samples except for one. This sample corresponded to a culture showing no growth whilst metagenomics identified *Aerococcus urinae* as being present (RA 88%, DNA yield 374 ng) (Figure 1 and Supplementary Data, Figure 3. The median difference in DI for groups UTI and non-UTI was significant (p<0.0001), with only 1 overlapping value between groups. The sensitivity and specificity of the DI for correctly allocating a urine sample to the UTI or non-UTI group were respectively 100% (95%-CI: 66.4-100%) and 96.7% (95%-CI: 83.8-99.9%)(23).

### Antimicrobial susceptibility

Phenotypic AST of the isolated uropathogens showed resistance against 0 to 6 drug classes per isolate, with resistance against an average of 3 drug classes per isolate, cumulating to 38 cases of AMR against a drug class for all 15 uropathogens together(Table 2, Supplementary data Table 3). Combined Resfinder and CARD analyses of WGS gave genotype-phenotype agreement in 22 of 38 resistant phenotypes.

Nine of 11 urine samples from the UTI group were available for metagenomic resistome analysis. These 9 samples contained 13 of the 15 cultured uropathogens, comprising 32 cases of resistance against an antimicrobial drug class, as per phenotypic AST. Metagenomic resistome analysis and phenotypic AST were concordant in 20 out of 32 cases (Table 2).

Five isolates (2 *E. cloacae,* 1 *K. pneumoniae*, 1 *M. morganii* and 1 *P. mirabilis*) derived from individual urine samples showed a nitrofurantoin resistance phenotype, but a nitrofurantoin resistance conferring gene was not detected, neither by metagenomic resistome nor by WGS analysis. Metagenomics did not detect a matching AMR gene to explain seven additional resistance phenotypes in five samples (Table 2) whilst AMR genes corresponding to these phenotypes were detected in the WGS assembly of the isolates.

In four samples, different AMR genes were identified by metagenomics and WGS of the corresponding isolate, although both could explain the phenotype (Table 2). In 2 of these samples (*bla*_SHV_ and *bla*_TEM_ genes detected by WGS in sample 19 and *bla*_DHA_ genes in sample 45), coverage of the metagenomic datasets was too low to distinguish the highly homologous gene variants (5-10 SNP differences between identified genes) (Supplementary data, Figure 4). The third, a *bla*_ACT-28_ gene detected by WGS in sample 7 was absent in the CosmosID database and was instead identified as the *bla*_ACT-9_ gene (a close variant with 5 SNPs difference) in the corresponding metagenomic sample. Finally, in sample 30, containing *E. coli* phenotypically resistant to 3^rd^ generation cephalosporins, metagenomic analysis identified an *ampC* gene whilst the *bla*_CTX-M-27_ ESBL-gene was identified in the WGS of the isolate.

Metagenomics outperformed WGS in three cases. In sample 28, containing *E. coli* phenotypically resistant to 3^rd^ generation cephalosporins, metagenomics identified an *ampC* gene, whilst WGS detected *bla*_*TEM-1B*_, which encodes resistance against penicillins, but not 3^rd^ generation cephalosporins. In two instances a resistance encoding genotype was identified by metagenomics but an equivalent coding gene was not detected in the corresponding WGS analysis. Both genotypes matched to a resistant phenotype: *oqxA* and *oqxB* coding for norfloxacin resistance in sample 45 and *ampC* for amoxicillin resistance in sample 23.

### Cost and timing

The time from urine arrival at the laboratory to obtaining results by culture was approximately 24h for non-UTI samples and 48-72h for culture and AST for UTI samples. The cost of this analysis per sample was approximately €4 for a non-UTI and €10 for a urine sample from the UTI group, excluding cost of staffing.

Total time required for metagenomic analysis was approximately 31h per sample, which included 4.5h for sample centrifugation, DNA extraction, and DNA shearing. A total of 15.5h were required for library preparation. Four hours were needed for sequencing. The computation time for CosmosID analysis for all 41 datasets was 3.88 min per sample using a Linux analysis server with two physical processors, a total of 12 cores (non-multithreaded) and 64GB of memory. The cost of library preparation and sequencing was €116 per sample, excluding cost of staffing. Should a screening be made based on quantity of extracted DNA per sample, non-UTI samples would have a processing time of approximately 4h and a cost of €5, namely the cost of DNA extraction only.

## DISCUSSION

This study demonstrates the potential of metagenomics for rapid detection of clinically relevant levels of uropathogens, thus distinguishing UTI from non-UTI urine samples.

Whilst several reports were previously published on the microbial community composition of urine(24, 25), successful metagenomic identification of uropathogens present in clinical samples has only been reported once to the best of our knowledge(7). However, that study did not include the semi-quantitative analysis required to establish a diagnosis of UTI(2, 3, 26). If metagenomics is to replace culture, a semi-quantitative analysis of clinically relevant bacteria present within the urine microbial community is critical to obtain a diagnostic test result, as has been done in this study.

Whilst all samples of the UTI group yielded >203.5 ng of DNA, 12 of 21 samples above this threshold were non-UTI urine samples, reflective of the fact that the procedure of urine sample collection is highly prone to bacterial contamination and that urine itself is not sterile(27). However, for none of the UTI samples was a low amount of DNA (<200 ng) associated with clinically relevant bacterial growth in culture, independent of the volume of urine from which DNA was extracted which ranged between 2 and 20 ml (data not shown). This observation suggests that establishing an unambiguous threshold of DNA extraction yield per urine volume is feasible due to the fact that in the presence of a UTI the bacterial DNA load in the urine increases exponentially. Thus, DNA quantity alone could serve as an initial screening step for a diagnostic algorithm, directing sequencing only those samples with a high DNA yield.

The total amount of time needed to detect and identify uropathogens and AMR genes in the UTI samples was significantly less (31h vs. 48-72hs) for metagenomics than for standard culture whilst metagenomics provided greater detail. Automated library preparation will further reduce processing time of UTI samples by several hours. Metagenomic costs were €116 vs. €10 for culture per sample. Although metagenomic sequencing is more expensive at the present time, sequencing costs are continuing to decline(28). At present €4 and 24h are required for ruling out a UTI by culture. Should a validated metagenomic diagnostic algorithm be in place, a UTI could be ruled out in a low yield DNA sample within 4.5 hours and costs would be reduced to the cost of DNA extraction – approximately €5 – making metagenomics highly competitive.

Overlapping values for the UTI and non-UTI samples were observed for both DNA quantity and relative abundance of uropathogens, indicating these parameters are unsuitable for identifying urine samples from patients with UTI. However, the DI discriminated UTI and non-UTI samples extremely well. The single outlier was a sample containing *Aerococcus urinae* in high relative abundance and high DNA yield, that had been reported to be culture negative, even though Gram-staining revealed Gram-positive cocci. *A. urinae* can be overlooked in cultures because of its morphological similarity to coagulase negative staphylococci(29), leading to false negative culture results.

Metagenomic analysis was successfully employed to identify uropathogens in all but two of the culture positive urine samples (Table 1). In the initial analysis *Morganella morganii* in sample 45 was not identified. The species was successfully identified after addition of the genome to CosmosID’s reference database, highlighting the importance of well-curated and complete databases. In sample 30, *S. aureus* was cultured at a concentration of >10^4^ CFU/ml. It is unlikely that insufficient DNA extraction due to poor lysis of Gram-positive bacteria can explain the apparent lack of *S. aureus* DNA, since an extensive pre-lysis protocol was used, although this cannot be ruled out(30). Alternatively, contamination of the agar plates may have occurred.

CosmosID metagenomic bioinformatics system identified AMR encoding genes that were confirmed to be present by pure isolate WGS analysis. However for a selected number of cases, metagenomic and WGS analyses identified a highly similar, but non-identical AMR gene. An explanation for this discrepancy can be the different databases used in the metagenomic and WGS analysis. For WGS, it has been shown that different bioinformatic AMR identification tools produce different results and that databases are often incomplete(31). In addition, low coverage of gene segments by metagenomic reads resulting in less accurate gene calling could explain differences between metagenomics and WGS. Interestingly, even though metagenomic and WGS methods identified different gene variants in those few instances, the variants were predicted to encode the same phenotype. However, accurate phenotype prediction on the basis of genomic information, whether WGS or metagenomics, remains a challenge. Phenotype clearly depends on more than the presence or absence of a resistance encoding gene. Whether antibiotic resistance mechanisms are expressed, depends on complex interplay between accumulating mutations or acquisition of resistance encoding genes and as yet unknown regulators, as well as environmental factors(32).

In conclusion, we provided a proof of concept of semi-quantitative metagenomic diagnostics for UTI, including the development of a “diagnostic index” based on uropathogen relative abundance and total DNA yield per sample, to facilitate rapid classification of urine samples to UTI and non-UTI groups. In future studies, clinical characteristics of patients and prior antibiotic use should be included to determine the sensitivity, specificity, and the positive and negative predictive value of this metagenomics analysis approach applied to urine samples using culture as the reference test.

## ACKNOWLEDGEMENTS

The authors would like to thank Cosimo Cristella for assistance in using RStudio and Frank Baas for critical input in sequencing planning and experiment design.

This study was supported by the COMPARE Consortium, which has received funding from the European Union’s Horizon 2020 research and innovation program under grant agreement No. 64347. The funders had no role in study design, data collection and interpretation, or the decision to submit the work for publication.

Rita R. Colwell is Founder and Chairman of the Board of CosmosID®, a bioinformatics company, and some of the other authors are employees of the company. Affiliation with CosmosID does not alter the authors’ adherence to all JCM policies as detailed in the online instructions for authors. Authors declare no conflict of interest with regards to this manuscript.

## AUTHORS CONTRIBUTION

V. A. J. wrote the manuscript. V. A. J., S. M. and M. E. J. performed the experiments. V. A. J., S. M., N. W., P. S. and N. A. H. performed the bio-informatics analysis. C. E. V. supervised the clinical diagnostic analysis. S. M., V. A. J., C. E. V., B. d W., R. R. C., M. D. d J., and C. S. designed the study. All authors provided critical review of the data and manuscript.

## References

1. Linder JA, Huang ES, Steinman MA, Gonzales R, Stafford RS. 2005. Fluoroquinolone prescribing in the United States: 1995 to 2002. Am J Med 118:259–68.

2. Hooton TM, Bradley SF, Cardenas DD, Colgan R, Geerlings SE, Rice JC, Saint S, Schaeffer AJ, Tambayh PA, Tenke P, Nicolle LE, Infectious Diseases Society of A. 2010. Diagnosis, prevention, and treatment of catheter-associated urinary tract infection in adults: 2009 International Clinical Practice Guidelines from the Infectious Diseases Society of America. Clin Infect Dis 50:625–63.

3. Gupta K, Hooton TM, Naber KG, Wullt B, Colgan R, Miller LG, Moran GJ, Nicolle LE, Raz R, Schaeffer AJ, Soper DE, Infectious Diseases Society of A, European Society for M, Infectious D. 2011. International clinical practice guidelines for the treatment of acute uncomplicated cystitis and pyelonephritis in women: A 2010 update by the Infectious Diseases Society of America and the European Society for Microbiology and Infectious Diseases. Clin Infect Dis 52:e103–20.

4. Cagnacci S, Gualco L, Debbia E, Schito GC, Marchese A. 2008. European emergence of ciprofloxacin-resistant Escherichia coli clonal groups O25:H4-ST 131 and O15:K52:H1 causing community-acquired uncomplicated cystitis. J Clin Microbiol 46:2605–12.

5. van der Zee A, Roorda L, Bosman G, Ossewaarde JM. 2016. Molecular Diagnosis of Urinary Tract Infections by Semi-Quantitative Detection of Uropathogens in a Routine Clinical Hospital Setting. PLoS One 11:e0150755.

6. Hansen WL, van der Donk CF, Bruggeman CA, Stobberingh EE, Wolffs PF. 2013. A real-time PCR-based semi-quantitative breakpoint to aid in molecular identification of urinary tract infections. PLoS One 8:e61439.

7. Hasman H, Saputra D, Sicheritz-Ponten T, Lund O, Svendsen CA, Frimodt-Moller N, Aarestrup FM. 2014. Rapid whole-genome sequencing for detection and characterization of microorganisms directly from clinical samples. J Clin Microbiol 52:139–46.

8. Hooton TM, Bradley SF, Cardenas DD, Colgan R, Geerlings SE, Rice JC, Saint S, Schaeffer AJ, Tambayh PA, Tenke P, Nicolle LE. 2010. Diagnosis, prevention, and treatment of catheter-associated urinary tract infection in adults: 2009 International Clinical Practice Guidelines from the Infectious Diseases Society of America. Clin Infect Dis 50:625–63.

9. Nicolle LE, Bradley S, Colgan R, Rice JC, Schaeffer A, Hooton TM. 2005. Infectious Diseases Society of America guidelines for the diagnosis and treatment of asymptomatic bacteriuria in adults. Clin Infect Dis 40:643–54.

10. EUCAST. Breakpoint tables for interpretation of MICs and zone diameters. Version 7.0, 2017. http://www.eucast.org Accessed 05-01-2017.

11. Bolger AM, Lohse M, Usadel B. 2014. Trimmomatic: a flexible trimmer for Illumina sequence data. Bioinformatics 30:2114–20.

12. Schmieder R, Edwards R. 2011. Fast identification and removal of sequence contamination from genomic and metagenomic datasets. PLoS One 6:e17288.

13. Ottesen A, Ramachandran P, Reed E, White JR, Hasan N, Subramanian P, Ryan G, Jarvis K, Grim C, Daquiqan N, Hanes D, Allard M, Colwell R, Brown E, Chen Y. 2016. Enrichment dynamics of Listeria monocytogenes and the associated microbiome from naturally contaminated ice cream linked to a listeriosis outbreak. BMC Microbiol 16:275.

14. Ponnusamy D, Kozlova EV, Sha J, Erova TE, Azar SR, Fitts EC, Kirtley ML, Tiner BL, Andersson JA, Grim CJ, Isom RP, Hasan NA, Colwell RR, Chopra AK. 2016. Cross-talk among flesh-eating Aeromonas hydrophila strains in mixed infection leading to necrotizing fasciitis. Proc Natl Acad Sci U S A 113:722–7.

15. Hasan NA, Young BA, Minard-Smith AT, Saeed K, Li H, Heizer EM, McMillan NJ, Isom R, Abdullah AS, Bornman DM, Faith SA, Choi SY, Dickens ML, Cebula TA, Colwell RR. 2014. Microbial community profiling of human saliva using shotgun metagenomic sequencing. PLoS One 9:e97699.

16. Lax S, Smith DP, Hampton-Marcell J, Owens SM, Handley KM, Scott NM, Gibbons SM, Larsen P, Shogan BD, Weiss S, Metcalf JL, Ursell LK, Vazquez-Baeza Y, Van Treuren W, Hasan NA, Gibson MK, Colwell R, Dantas G, Knight R, Gilbert JA. 2014. Longitudinal analysis of microbial interaction between humans and the indoor environment. Science 345:1048–52.

17. Foxman B. 2010. The epidemiology of urinary tract infection. Nat Rev Urol 7:653–60.

18. Bankevich A, Nurk S, Antipov D, Gurevich AA, Dvorkin M, Kulikov AS, Lesin VM, Nikolenko SI, Pham S, Prjibelski AD, Pyshkin AV, Sirotkin AV, Vyahhi N, Tesler G, Alekseyev MA, Pevzner PA. 2012. SPAdes: a new genome assembly algorithm and its applications to single-cell sequencing. J Comput Biol 19:455–77.

19. Larsen MV, Cosentino S, Lukjancenko O, Saputra D, Rasmussen S, Hasman H, Sicheritz-Ponten T, Aarestrup FM, Ussery DW, Lund O. 2014. Benchmarking of methods for genomic taxonomy. J Clin Microbiol 52:1529–39.

20. Mount DW. 2007. Using the Basic Local Alignment Search Tool (BLAST). CSH Protoc 2007:pdb top17.

21. Zankari E, Hasman H, Cosentino S, Vestergaard M, Rasmussen S, Lund O, Aarestrup FM, Larsen MV. 2012. Identification of acquired antimicrobial resistance genes. J Antimicrob Chemother 67:2640–4.

22. Jia B, Raphenya AR, Alcock B, Waglechner N, Guo P, Tsang KK, Lago BA, Dave BM, Pereira S, Sharma AN, Doshi S, Courtot M, Lo R, Williams LE, Frye JG, Elsayegh T, Sardar D, Westman EL, Pawlowski AC, Johnson TA, Brinkman FS, Wright GD, McArthur AG. 2017. CARD 2017: expansion and model–centric curation of the comprehensive antibiotic resistance database. Nucleic Acids Res 45:D566–D573.

23. Griner PF, Mayewski RJ, Mushlin AI, Greenland P. 1981. Selection and interpretation of diagnostic tests and procedures. Principles and applications. Ann Intern Med 94:557–92.

24. Hilt EE, McKinley K, Pearce MM, Rosenfeld AB, Zilliox MJ, Mueller ER, Brubaker L, Gai X, Wolfe AJ, Schreckenberger PC. 2014. Urine Is Not Sterile: Use of Enhanced Urine Culture Techniques To Detect Resident Bacterial Flora in the Adult Female Bladder. Journal of Clinical Microbiology 52:871–876.

25. Siddiqui H, Lagesen K, Nederbragt AJ, Jeansson SL, Jakobsen KS. 2012. Alterations of microbiota in urine from women with interstitial cystitis. BMC Microbiology 12:205.

26. Burd EM, Kehl KS. 2011. A Critical Appraisal of the Role of the Clinical Microbiology Laboratory in the Diagnosis of Urinary Tract Infections. Journal of Clinical Microbiology 49:S34–S38.

27. Thomas-White K, Brady M, Wolfe AJ, Mueller ER. 2016. The bladder is not sterile: History and current discoveries on the urinary microbiome. Curr Bladder Dysfunct Rep 11:18–24.

28. NIH NHGRI. July 6, 2016 2016. https://www.genome.gov/sequencingcosts/. Accessed June 8, 2017.

29. Grude N, Jenkins A, Tveten Y, Kristiansen BE. 2003. Identification of Aerococcus urinae in urine samples. Clinical Microbiology and Infection 9:976–979.

30. Ezaki T, Suzuki S. 1982. Achromopeptidase for lysis of anaerobic gram-positive cocci. Journal of Clinical Microbiology 16:844–846.

31. Xavier BB, Das AJ, Cochrane G, De Ganck S, Kumar-Singh S, Aarestrup FM, Goossens H, Malhotra-Kumar S. 2016. Consolidating and exploring antibiotic resistance gene data resources. Journal of Clinical Microbiology doi:10.1128/jcm.02717-15.

32. Hughes D, Andersson DI. 2017. Environmental and genetic modulation of the phenotypic expression of antibiotic resistance. FEMS Microbiology Reviews 41:374–391.

